# Cyclosporine A inhibits MRTF-SRF signalling through Na^+^/K^+^ ATPase inhibition & Actin remodelling

**DOI:** 10.1101/492843

**Authors:** Bastien Burat, Quentin Faucher, Petra Čechová, Hélène Arnion, Florent Di Meo, François-Ludovic Sauvage, Pierre Marquet, Marie Essig

## Abstract

Calcineurin Inhibitors (CNI) are the pillars of immunosuppression in transplantation. However, they display a potent nephrotoxicity whose mechanisms remained widely unsolved. We used an untargeted quantitative proteomic approach (iTRAQ technology) to highlight new targets of CNI in renal proximal tubular cells (RPTCs). CNI-treated RPTCs proteome displayed an over-representation of Actin-binding proteins with a CNI-specific expression profile. Cyclosporine A (CsA) induced F-Actin remodelling and depolymerisation, decreased F-Actin-stabilizing, polymerization-promoting Cofilin (CFL) oligomers and inhibited the G-Actin-regulated serum responsive factor (SRF) pathway. Inhibition of CFL canonical phosphorylation pathway reproduced CsA effects; however, Ser3, an analogue of the phosphorylation site of CFL prevented the effects of CsA which suggests that CsA acted independently from the canonical CFL regulation. CFL is known to be regulated by the Na^+^/K^+^-ATPase. Molecular docking calculations evidenced 2 inhibiting sites of CsA on Na^+^/K^+^-ATPase and a 23% decrease in Na^+^/K^+^-ATPase activity of RPTCs was observed with CsA. Ouabain, a specific inhibitor of Na^+^/K^+^-ATPase also reproduced CsA effects on Actin organization and SRF activity. Altogether, these results described a new original pathway explaining CsA nephrotoxicity.

## Introduction

The development of immunosuppressive regimens based on the Calcineurin Inhibitors (CNI), namely Cyclosporine A (CsA) and Tacrolimus (Tac), has been a breakthrough in the prevention of allograft rejection in solid organ transplantation (Calne *et al*, 1978,1979; Starzl *et al*, 1989). Although CNI are now widely used in clinical protocols of immunosuppression (Hart *et al*, 2016, 2017) and have significantly improved short-term graft and patient survival, long-term exposure is associated with major limiting deleterious side effects such as nephrotoxicity, leading to end-stage renal disease (Myers *et al*, 1984).

CNI nephrotoxicity evolves from acute, reversible vascular and hemodynamic impairments to chronic irreversible and generalized renal injuries (Gaston, 2009; Naesens *et al*, 2009). In particular, chronic CNI nephrotoxicity results in interstitial fibrosis and tubular atrophy (IF/TA) of proximal tubules, whose mechanisms remain largely unsolved. Up to now, CNI are known to : disrupt cell cycle and induce cell death (Lally *et al*, 1999; Jennings *et al*, 2007; Ito *et al*, 1995; Healy *et al*, 1998; Ortiz *et al*, 1998; Justo, 2003); induce endoplasmic reticulum stress and unfolded protein response (Pallet *et al*, 2008a, 2008c; Han *et al*, 2008; Hama *et al*, 2013; Pallet *et al*, 2008b); promote oxidative stress (Vetter *et al*, 2003; Djamali, 2007); impact ion homeostasis (Heering & Grabensee, 1991), or induce epithelial-mesenchymal transition (EMT) (Hazzan *et al*, 2011; McMorrow *et al*, 2005; Slattery *et al*, 2005).

These observations resulting from targeted experimental approaches only partially described CNI cell toxicity and do not allow a complete understanding of all the pathophysiological mechanisms at stake. Because the mechanisms of CNI side effects remain widely unsolved by targeted strategies, the design of untargeted experiments is of utmost importance to gain new insights and improve knowledge about the pathophysiology of CNI proximal tubule cell (PTCs) toxicity.

Omics based on the high-throughput analysis of biological systems such as Shotgun proteomics seem particularly well suited to this purpose. In the present study, we performed the quantitative proteomic analysis and dynamic mapping of CNI-exposed PTC proteome to elucidate new intracellular pathways specifically modified under CNI exposure.

Upon the significant over-representation and differential expression of Actin family cytoskeletal proteins, we focused on the deciphering of the intracellular mechanisms of CsA-induced reorganization of the Actin cytoskeleton of PTC and its downstream consequences. We showed that CsA induced an inhibition of the Actin-dependent Myocardin-Related Transcription Factors-Serum Response Factor (MRTF-SRF) transcription activity through an original regulation of Cofilin (CFL) by the Na^+^/K^+^-ATPase.

## Results

### Quantitative proteomic analysis of PTC proteome & mapping of CNi-induced perturbations highlight the over-representation and differential expression of Actin family cytoskeletal proteins

iTRAQ technology allowed the identification of 130 proteins in PTC lysate. The PANTHER over-representation test (Table 1) classified 128 *Sus scrofa* proteins, related to ribosome structure (PC00202, fold enrichment = 15.90, p-value = 1.48.10-18***), nucleic acid binding (PC00031, fold enrichment = 5.87, p-value = 6.08.10-10*** ; PC00171, fold enrichment = 2.30, p-value = 4.86.10-3**), metabolism processes (PC00120, fold enrichment = 31.24, p-value = 2.70.10-2* ; PC00092, fold enrichment = 7.83, p-value = 4.19.10-5*** ; PC00135, fold enrichment = 7.63, p-value = 3.09.10-2* ; PC00176, fold enrichment = 5.86, p-value = 1.46.10-7***) and cytoskeleton dynamics (PC00228, fold enrichment = 33.32, p-value = 1.51.10-3** ; PC00041, fold enrichment = 5.32, p-value = 4.39.10-3** ; PC00085, fold enrichment = 3.79, p-value = 2.24.10-3**).

**Table 1.**
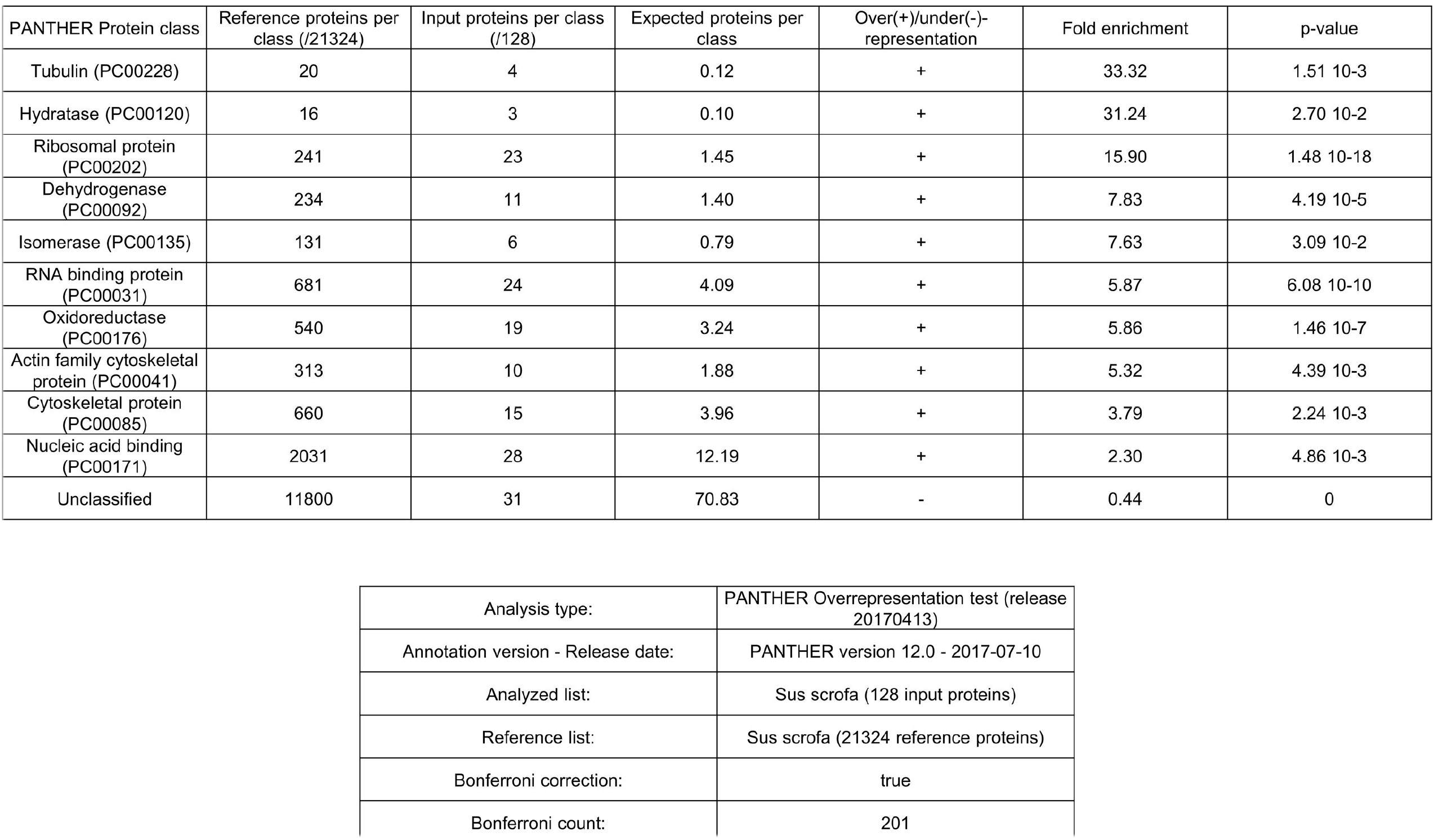
Cytoskeletal proteins are significantly overrepresented in the proteome of CNI-exposed LLC PK-1 monitored by iTRAQ shotgun proteomics. Swiss-Prot IDs from 128 identified proteins were submitted to the PANTHER Overrepresentation test (release date 20170413) parsing the PANTHER database (version 12.0, released on 2017-07-10) using the *Sus scrofa* reference list and the PANTHER Protein Class annotation data set. Bonferroni correction was applied.

Among the 15 proteins related to cytoskeleton structure and dynamics, Actin family cytoskeletal proteins (PANTHER Protein Class PC00041, fold enrichment = 5.32, p-value = 4.39.10-3**) formed a node of 10 interconnected proteins in the visualisation of functional protein networks (Figure EV1 A).

Among the 70 proteins with significant variations of abundance, Actin family cytoskeletal proteins like Moesin, Radixin, Myosin VI, α-SMA, Troponin-C, F-Actin capping protein subunits α2 and β, and Cofilin-1 showed a CNI-specific expression profile (Figure EV1 B).

### CsA, but not Tac, elicited a strong reorganization of Actin cytoskeleton of PTCs

In fluorescence microscopy, cell monolayers of proximal tubular LLC PK-1 showcased a puzzled pattern as phalloidin-labelled F-Actin cytoskeleton organized cell membranes into intercellular convolutions (Figure 1 A, panel a). These peculiar structuring of plasma membranes were reminiscent of the way basolateral membranes connect inside proximal tubular epithelia, in vivo.

**Figure 1.**
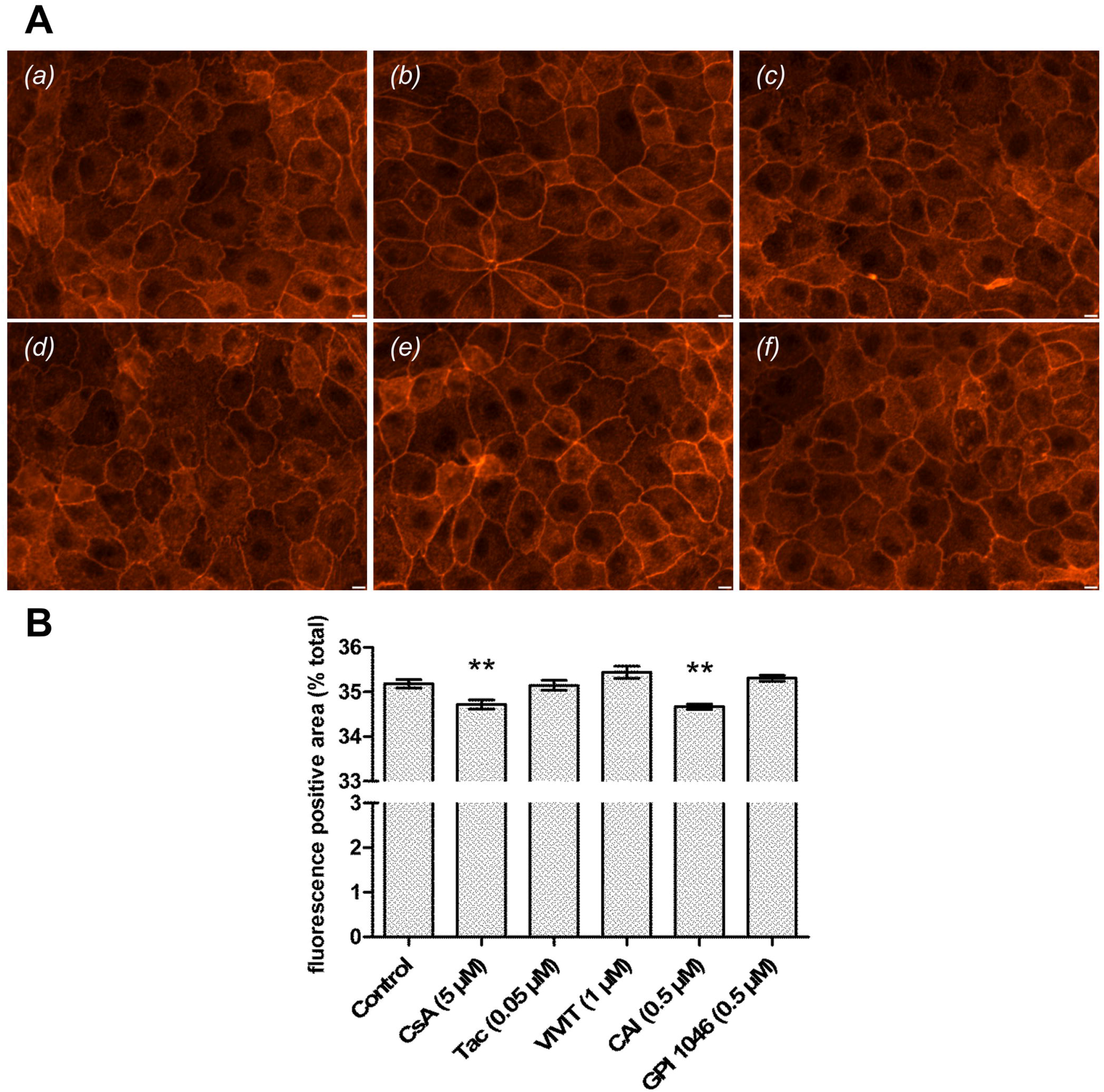
CsA, but not TAC, triggers F-Actin depolymerization in a NFAT-independent, immunophilin inhibition-related way. A. Fluorescence images of LLC PK-1 Actin cytoskeleton labelled with TRITC-Phalloidin. Scale bar 10 μm
B. Quantification of red fluorescence-positive area. Mean ± SEM. One-way ANOVA plus Dunnett’s post-test (** p<0.01) Drug condition: (a) Vehicle (0.5 % Ethanol) (b) 5 μM CsA (c) 0.05 μM Tac (d) 1 μM VIVIT (e) 0.5 μM CAI (f) 0.5 μM GPI 1046. Exposure time: 24 h (n=3).

Upon 24-hour CsA exposure, LLC PK-1 sustained a significant reorganization of cortical Actin cytoskeleton (Figure 1 A, panel b) related to a significant loss of F-Actin-based structures (Figure 1 B, -0.46 %, p<0.01**). CsA effects seemed to be related to the inhibition of the peptidylprolyl cis-trans isomerase activity of its target immunophilin Cyclophilin A (CypA) since a specific pharmacological inhibitor of CypA (CAI), without any immunological effect, elicited similar Actin modifications (Figure 1 A panel e, Figure 1 B, -0.51 %, p<0.01**). On the contrary, CsA effects seemed independent of Calcineurin inhibition since neither Tac, VIVIT (a specific inhibitor of NFAT dephosphorylation by Calcineurin) nor GPI 1046 (a pharmacological inhibitor of FKBP12) exhibited modifications of the Actin organization (Figure 1A panels c, d and f, Figure 1B).

In conclusion, these observations suggested that CsA triggered the deep remodelling of proximal tubular Actin cytoskeleton and the depolymerization of F-Actin, in a Cyclophilin-dependent way, independently of Calcineurin inhibition.

### Actin depolymerization led to a decrease in MRTF-SRF transcription activity in LLC PK-1 upon CsA exposure

Upon 24-hour CsA exposure, SRF transcription activity was significantly reduced (Figure 2 A, -48.5 % of control activity, p<0.001***). Likewise, CAI elicited a concentration-dependent negative regulation of SRF (Figure 2 C). On the contrary, neither Tac, VIVIT (Figure 2 B) nor GPI 1046 (Figure 2 D) reduced SRF transcription activity which was slightly increased upon Tac and VIVIT exposure. Profiles of drug-related Actin dynamics strictly superimposed to profiles of drug-related SRF transcription activity.

**Figure 2.**
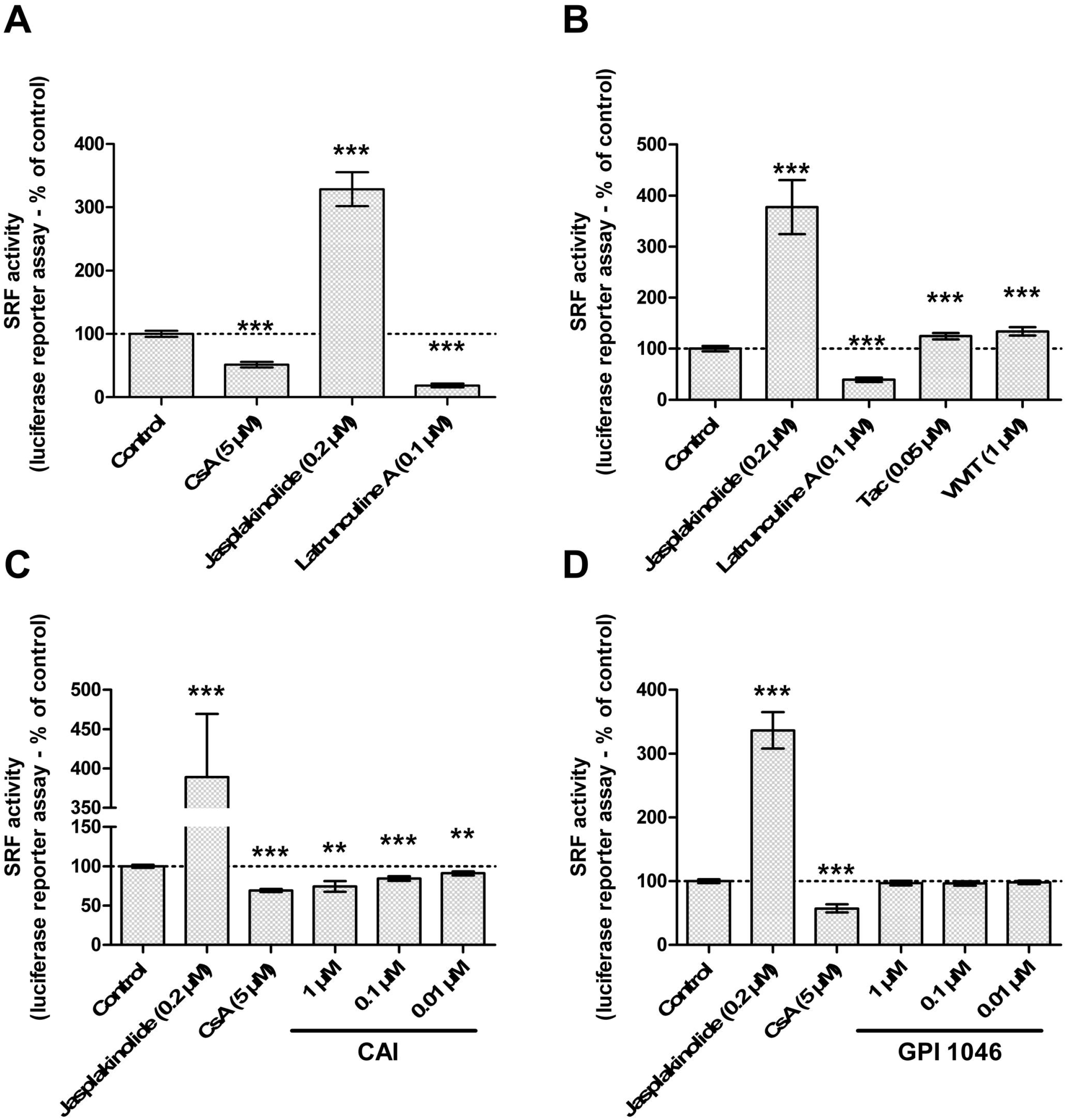
CsA-induced accumulation of G-Actin leads to inhibition of SRF-MRTF transcription activity in a NFAT-independent, immunophilin inhibition-related way. SRF transcription activity was measured by luciferase gene reporter assay in LLC PK-1 SRE exposed for 24 h to: Drug condition: A-D. Vehicle (0.5% Ethanol-0,2% DMSO), 5 μM CsA, 0.2 μM Jasplakinolide; A-B. 0.1 μM Latrunculin A; B. 0.05 μM Tac, 1 μM VIVIT; C. 0.01-1 μM CAI, D. 0.01-1 μM GPI 1046. Mean ± SEM. One-sample t-test (p<0.01**, p<0,001***). A-C, D (n=3). B (n=5).

In conclusion, CsA downregulated the transcription of MRTF-SRF target genes, in a Cyclophilin-dependent way, independently of Calcineurin inhibition.

### CsA regulates Actin-binding protein Cofilin through changes in Cofilin-Actin ratio and oligomerization state

Differential quantitative proteomic analysis of identified Actin family cytoskeletal proteins (PANTHER Protein Class PC00041) showed that CsA decreased the CFL: Actin ratio since Actin was overexpressed while CFL levels remained steady. On the contrary, Tac induced equivalent overexpression for both Actin and CFL (Figure EV1).

Western blot analysis of cross-linked CFL oligomers (Figure 3 A) indicated that CsA induced a significant decrease in dimers and tetramers (RdiCFL = 0.40 ± 0.09; RtetraCFL = 0.38 ± 0.16) while monomers remain steady (RmonoCFL = 0.90 ± 0.06) (Figure 3 B).

**Figure 3.**
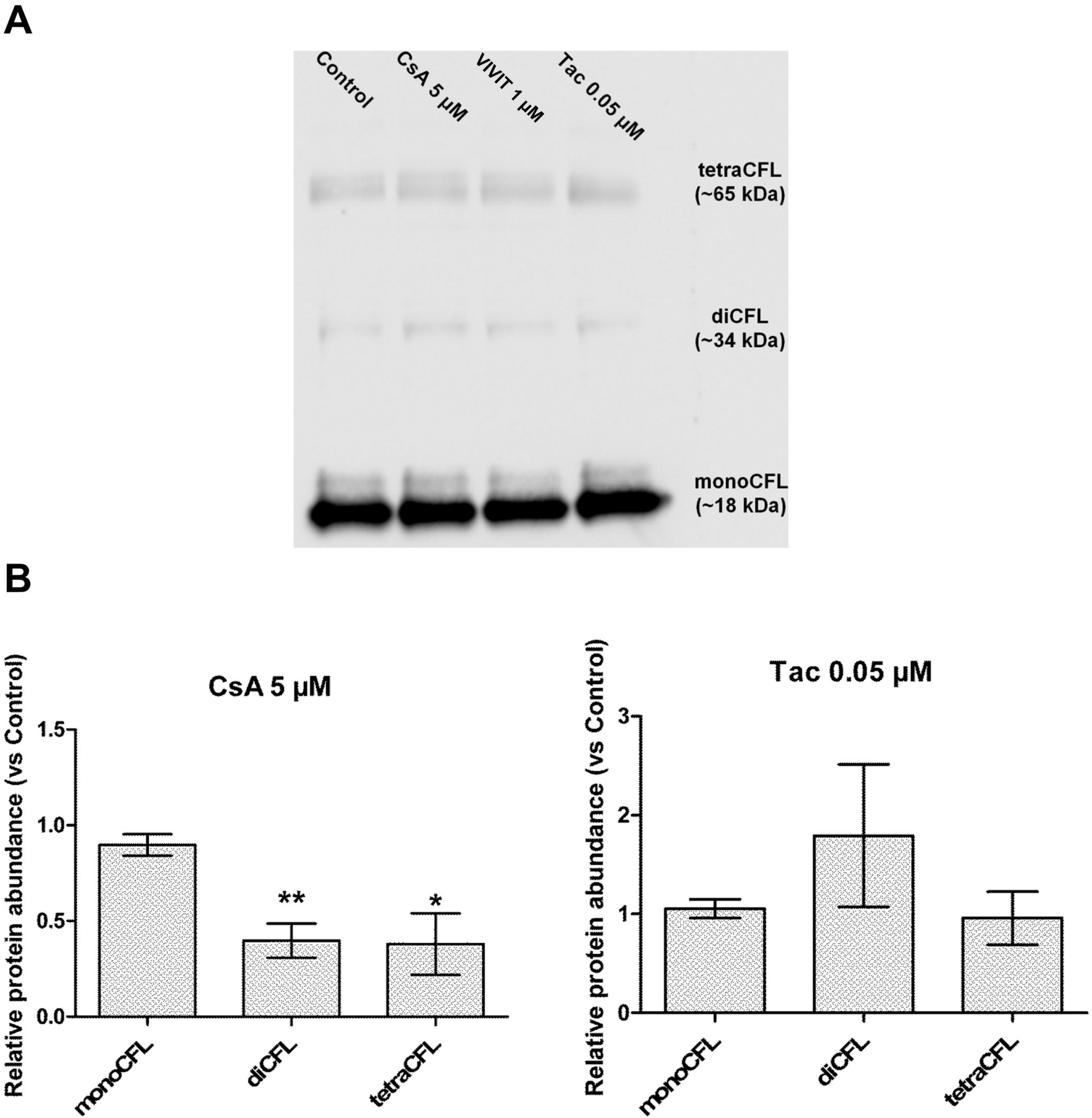
CsA, but not Tac, induced changes in CFL concentration-to-Actin A. Western blot detection of formaldehyde-cross-linked CFL oligomers in LLC PK-1 lysates.
B. Quantification of relative protein abundance of CFL oligomeric forms. Mean ± SEM. One-way ANOVA plus Dunnett’s post-test (*p<0.05, **p<0.01). Drug condition: 5 μM CsA, 0.05 μM Tac. Exposure time: 24 h (n=5)

In conclusion, CsA elicited a shift in the balance between oligomeric forms of CFL, decreasing the protein abundance of CFL dimers and tetramers.

### CsA effects seemed partly independent of CFL phosphorylation/dephosphorylation

The S3-R peptide, an analogue of the CFL phosphorylation site and a non-specific inhibitor of Ser3-targeting proteins, did not affect the organization of the Actin cytoskeleton (Figure 4 A, panel b) but was linked to a slight yet significant increase in F-Actin levels (Figure 4 B, +0.36 %, p<0.05*). In association with CsA, S3-R nullified the CsA-induced Actin remodelling (Figure 4 A, panel e, Figure 4 B). Conversely, the addition of a specific pharmacological inhibitor of LIMK, LIMKi3, alone or associated with CsA triggered CsA-like phenotype of Actin dynamics (Figure 4 A, panel c & f) and significantly decreased F-Actin levels (Figure 4 B, LIMKi3 alone: -1.62 %, p<0.001***; LIMKi3 + CsA: -1.22 %, p<0.001***).

**Figure 4.**
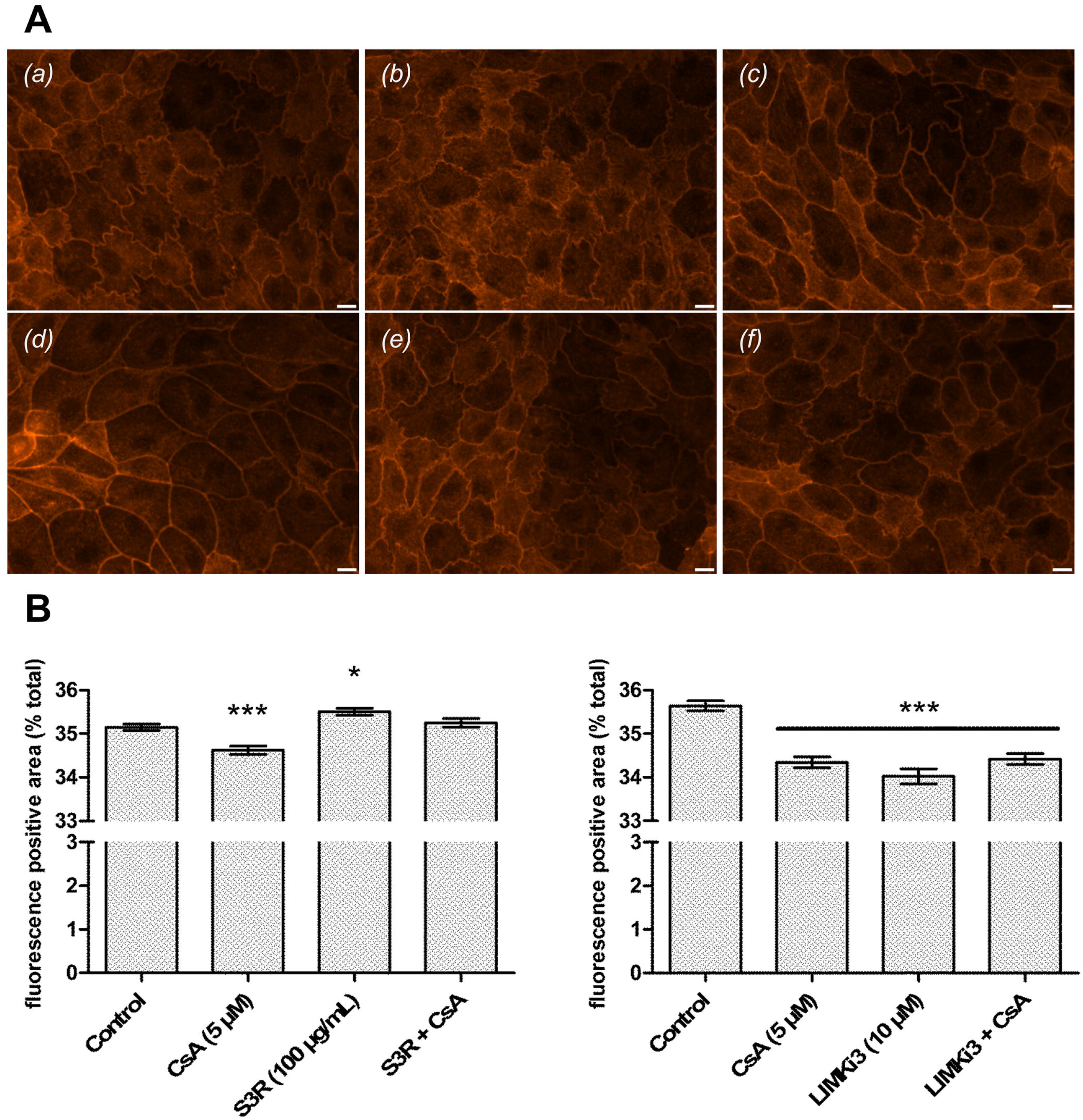
Cofilin is involved in CsA-induced Actin reorganization through its phosphorylation site A. Fluorescence images of LLC PK-1 Actin cytoskeleton labelled with TRITC-Phalloidin. Scale bar 10 μm
B. Quantification of red fluorescence-positive area. Mean ± SEM. One-way ANOVA plus Tukey’s post-test (* p<0.05,*** p<0.001) (n=4). Drug condition: (a) Vehicle (b) 100 μg/mL S3R (c) 10 μM LIMKi3 (d) 5 μM CsA (e) S3R + CsA (f) LIMKi3 + CsA. Exposure time: 24 h (n=4).

S3-R alone did not impact the MRTF-SRF transcription activity (Figure 5 A). In association with CsA, S3-R significantly decreased the CsA-induced inhibition of MRTF-SRF transcription activity (CsA: -33.1 %; S3-R + CsA: −24.5 %; +8.6 %, p<0.05*). LIMKi3 alone induced an inhibition of MRTF-SRF transcription activity (Figure 5 B, −52.3 %, p<0.001***) like what was observed with CsA alone (−35.9 %, p<0.001***). Furthermore, LIMKi3 reinforced the CsA-induced inhibition of SRF transcription activity (−61.5 %, p<0.001***).

**Figure 5.**
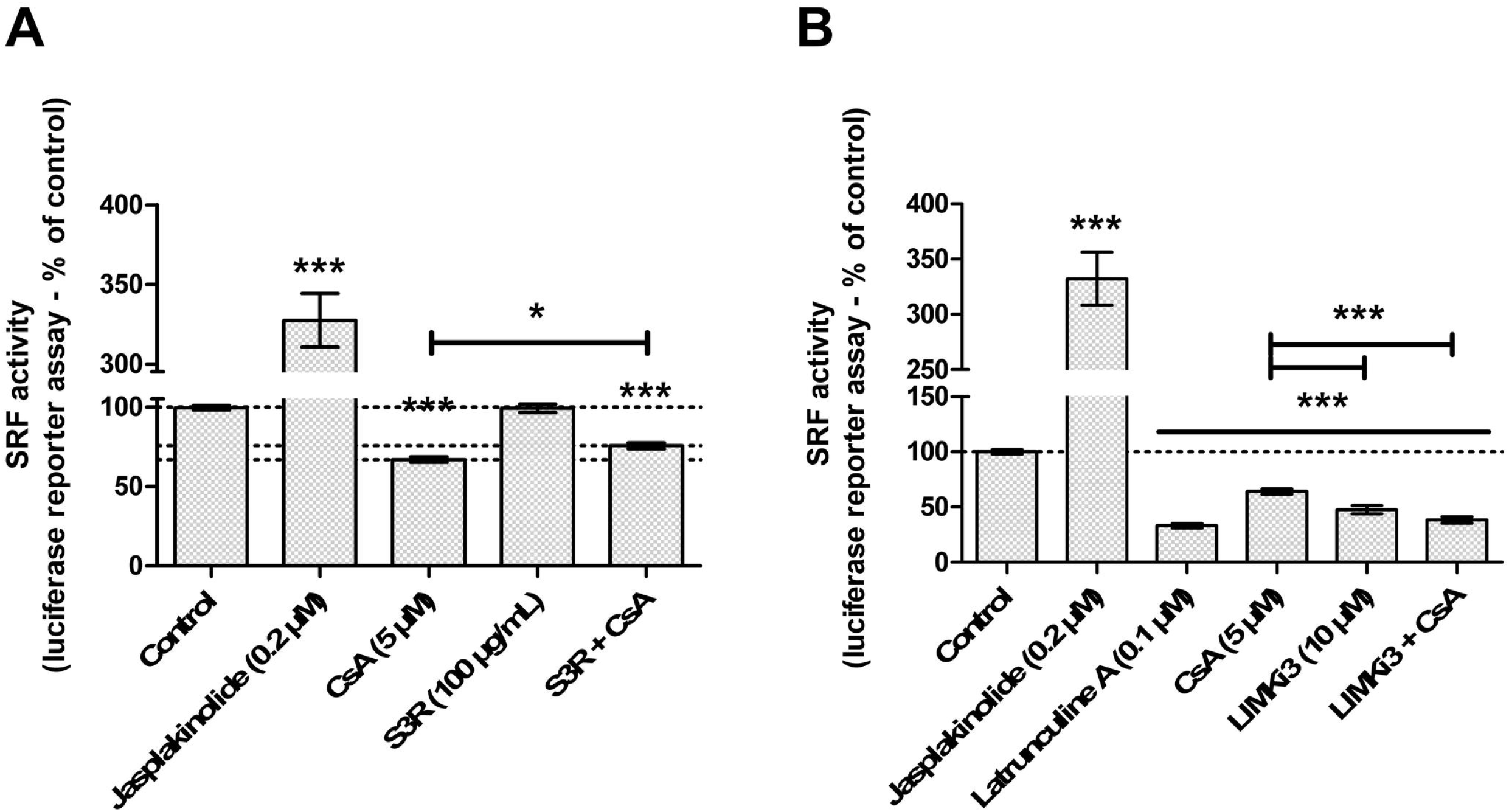
Cofilin is involved in CsA-induced inhibtion of MRTF-SRF transcription activity. SRF transcription activity was measured by luciferase gene reporter assay in LLC PK-1 SRE exposed 24 h to: A. S3-R 100 μg/mL; S3R + CsA. (n=6)
B. LIMKi3 10 μM ; LIMKi3 + CsA. (n=7) Mean ± SEM. One-sample t-test for versus control comparison, One-way ANOVA plus Tukey’s post-test for multiple condition comparison (p<0.01*, p<0.001***).

The pharmacological inhibitors of the RhoGTPases pathway, Y27632 (ROCK inhibitor) and EHT1864 (Rac1 inhibitor) elicited the same effects as LIMKi3 at both cytoskeletal (Figure EV2 A) and transcriptional (Figure EV2 B) levels.

In conclusion, CsA effects involved the phosphorylation site of CFL as a target for dephosphorylation. Inhibitors of the RhoGTPases pathway triggered CsA-like effects.

### CsA inhibited Na^+^/K^+^-ATPase activity in PTCs and Ouabain-induced inhibition of Na^+^/K^+^-ATPase mimicked Actin reorganization and inhibition of MRTF-SRF transcription activity observed upon CsA exposure

A significant decrease of the Na^+^/K^+^-ATPase activity (−23.1 % of control, p<0.01**) was observed in LLC PK-1 exposed for 24 h to CsA (Figure 6 A) whereas Tac treatment had no effect.

**Figure 6.**
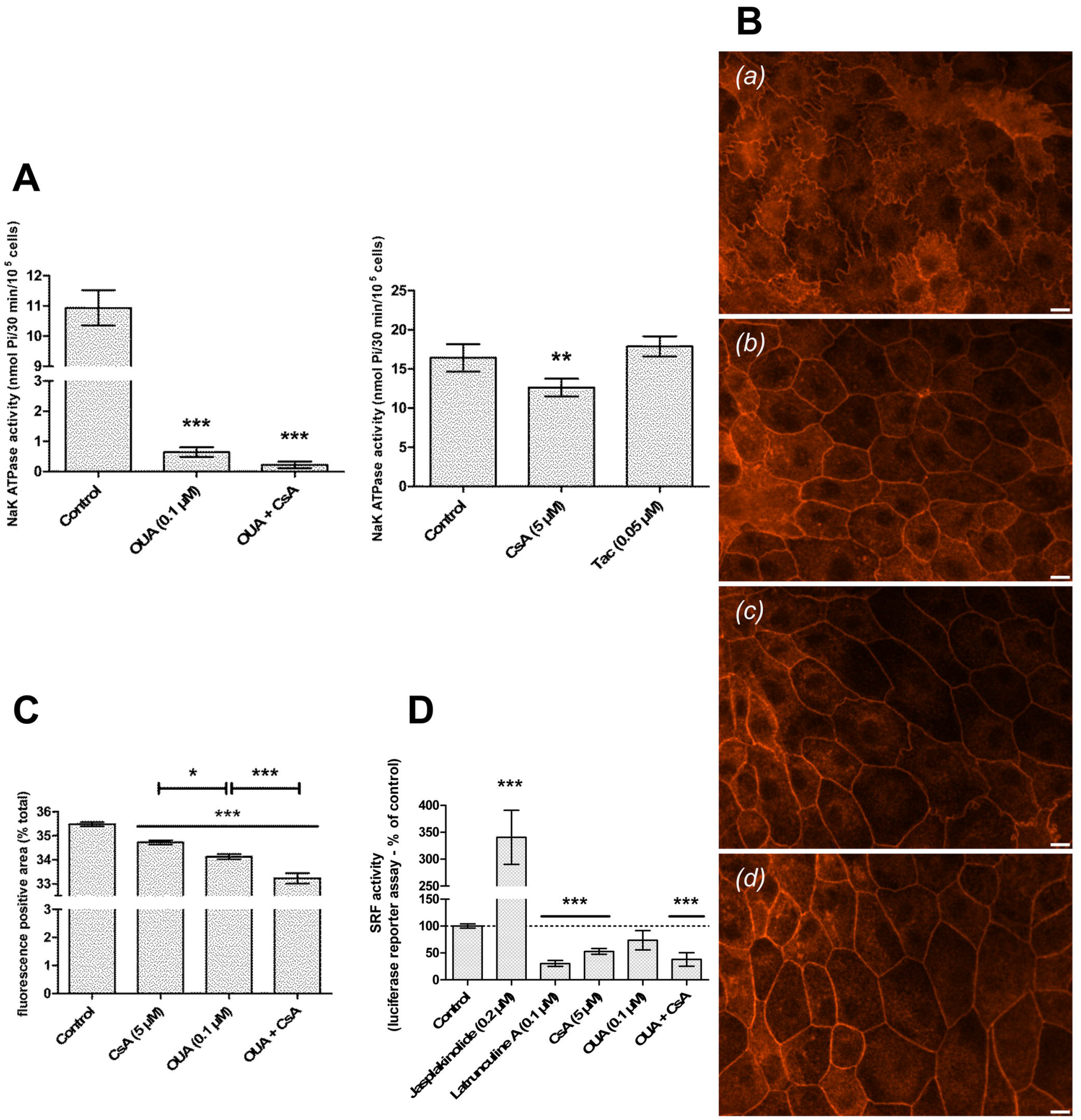
CsA partially inhibits Na^+^/K^+^-ATPase activity; Na^+^/K^+^-ATPase inhibitor Ouabain triggers and potentiates CsA-like actin reorganization and inhibition of SRF-MRTF transcription activity A. Na^+^/K^+^-ATPase activity was measured by colorimetric phosphate assay in LLC PK-1. Mean ± SEM. One-way ANOVA plus Dunn’s post-test (p<0.01**,p<0.001***) (n=3)
B. Fluorescence images of LLC PK-1 Actin cytoskeleton labelled with TRITC-Phalloidin. Scale bar 10 μm.
C. Quantification of red fluorescence-positive area. Mean ± SEM, One-way ANOVA plus Tukey’s post-test (p<0.01*, p<0.001***) (n=4).
D. SRF transcription activity was measured by luciferase gene reporter assay in LLC PK-1 SRE. One-sample t-test for versus control comparison, One-way ANOVA plus Tukey’s post-test for multiple condition comparison (p<0.001***) (n=4) Drug condition: (a) Vehicle (b) 5 μM CsA (c) 100 nM Ouabain (d) CsA + Ouabain. Exposure time: 24 h.

LLC PK-1 exposed for 24 h to 100 nM Ouabain featured CsA-like Actin reorganization (Figure 6 B panel c) with a significantly greater loss of F-Actin (Figure 6 C, CsA: -0.76 %; OUA: −1.35 %, p<0.001***). Moreover, Ouabain co-treatment with CsA potentiated CsA-induced cell stiffening (Figure 6 B panel d) and F-Actin decrease (Figure 6 C, −2.26 %, p<0.001***). Likewise, Ouabain exposure led to the inhibition of MRTF-SRF transcription activity (Figure 6 D, −26.6 %). Ouabain co-treatment with CsA reinforced CsA effects (CsA: – 47.0 %; OUA + CsA: −62.0 %, p<0.001***).

In conclusion, CsA targeted Na^+^/K^+^-ATPase and inhibited its activity. Na^+^/K^+^-ATPase inhibitor Ouabain elicited Actin reorganization the same way CsA did. Ouabain-related Actin dynamics induced the partial inhibition of MRTF-SRF transcription activity. The association of Ouabain with CsA tended to potentiate CsA effects on tubular proximal morphology and Actin dynamics.

### CsA molecular docking into Na^+^/K^+^-ATPase

CsA was docked into the closed (E1) conformation of Na^+^/K^+^-ATPase with and without ATP and into the open (E2) conformation without any ligands (Figure EV3). All the obtained binding poses had similar binding affinity, ranging from −7.7 to −6.6 kcal.mol-1. Assuming the known membrane partitioning of CsA (Haynes *et al*, 1985; Wang *et al*, 2018), only the binding poses close to the membrane boundary were considered.

On the open conformation of Na^+^/K^+^-ATPase (state E2, Figure EV3 A), there is a binding site at the C-terminal end of the protein, near the β-subunit, in the vicinity of the C-terminal pathway (Poulsen *et al*, 2010; Čechová *et al*, 2016). There is another binding site under the A-domain. It is noteworthy that this binding site is present in both conformations (Figure EV3 A and B) and near the binding site of Oligomycin A in the 3WGV crystal structure (Kanai *et al*, 2013).

In the closed conformation of Na^+^/K^+^-ATPase (state E1, Figure EV3 B and C), the most prominent binding sites are at the lipid-binding sites as previously defined (Cornelius *et al*, 2015). Most of the binding poses are located at the so-called site C – a position of bound cholesterol and phospholipid, involving aromatic and aliphatic residues such as W32 and Y39 or V838. This site has been previously implicated in the Na^+^/K^+^-ATPase inhibition.

## Discussion

In this work, we demonstrated that CsA-induced remodelling of the Actin cytoskeleton of PTC was associated with a modification of the F-Actin: G-Actin ratio through an original regulation of CFL: Indeed, CsA exposure decreased the total CFL: Actin ratio as well as the levels of CFL dimers and tetramers, independently of the phosphorylation by LIM kinases but most likely related to the CsA-induced inhibition of Na^+^/K^+^-ATPase.

In the kidney, the study of podocytes exposed to CsA revealed that the dynamic of the Actin cytoskeleton was sensitive to CsA. Indeed, CsA exerted the stabilization of the Actin cytoskeleton of podocyte foot processes, explaining the anti-proteinuric effect of CsA which is observed in nephrosis. This mechanism was demonstrated to be Calcineurin-dependent with the inhibition of the dephosphorylation and consequent proteolysis of Synaptopodin, a podocyte specific Actin-binding protein (ABP) whose activity is essential for podocyte structure-function (Faul *et al*, 2008). Besides, CsA induced the overexpression of CFL, a ubiquitary ABP, and its regulation in a non-phosphorylated state, independently from the effects on Synaptopodin (Li *et al*, 2014).

In the light of early observations of CsA effects on the Actin cytoskeleton of podocytes, the study of the Actin dynamic in PTC appeared of utmost interest in the elucidation of pathophysiological mechanisms of tubular atrophy upon CNI exposure. In PTC, we demonstrated that the Actin cytoskeleton could be modified by CsA (Descazeaud *et al*, 2012). Indeed, independently from the inhibition of NFAT, CsA triggered a reversible disorganization of Actin scaffolding at the periphery of the cell. Whether a mechanism involving the CsA-related regulation of CFL was responsible for the effects of CsA on PTC remained to be addressed.

CFL was initially described as an Actin-depolymerizing factor involved in Actin treadmilling. CFL modifies the physical-chemical properties of the microfilaments, promotes Pi release and ADP-Actin-G dissociation from older segments to replenish the pool of G-Actin and to supply enough material for microfilament renewal. Since then, the scope of CFL activity was expanded, in the light of the multiple mechanisms of CFL regulation. Up to date, CFL activity is known to be sensitive to F-Actin saturation, phosphorylation-dephosphorylation of its Ser3 residue, PIP2 interaction, intracellular pH and oxidative stress (Vantroys *et al*, 2008).

Since direct interaction with F-Actin microfilaments is necessary for the Actin-related activity CFL, the degree of saturation of the microfilament (or the density of CFL near the microfilament) play a major role in the regulation of CFL. Saturation is correlated to global CFL concentration, CFL: Actin ratio and CFL oligomerization. At high saturation and CFL density, F-actin filaments are severed and prone to the nucleation and polymerization of new branches. Conversely, at low saturation and CFL density, F-actin filaments are severed and completely depolymerized into G-Actin. Low CFL concentrations are related to low saturation, and so, the higher the concentration, the higher the saturation of microfilaments (Andrianantoandro & Pollard, 2006; Yeoh *et al*, 2002). Besides, in vivo and in vitro, CFL may exist as monomers, dimers and tetramers with higher-order oligomers related to higher CFL concentration and F-Actin saturation (Pfannstiel *et al*, 2001; Goyal *et al*, 2013).

The switches between phosphorylated and dephosphorylated status are an-other level of regulation between active and inactive CFL. Phosphorylation of the Ser3 residue prevents CFL-Actin interaction while dephosphorylation rescues CFL-Actin interaction and enables Actin-related CFL activity. Phosphorylation of Ser3 is only catalysed by LIM kinases, downstream small RhoGTPases and their associated kinases (Geneste *et al*, 2002). Dephosphorylation of CFL Ser3 is catalysed by Slingshot phosphatases (Huang *et al*, 2006), which itself is activated by Calcineurin.

The regulation of CFL orientates Actin dynamics towards the polymerization of G-Actin monomers into F-Actin microfilaments, or, conversely, promotes the depolymerization of F-actin into G-Actin. In case of a cytoplasmic accumulation of G-Actin, G-Actin traps Myocardin-Related Transcription Factor (MRTF), a co-factor of the Serum Response Factor (SRF) transcription factor. Complexes of MRTF-G-Actin are restricted to the cytoplasm. Gene transcription by MRTF-SRF complexes depends on the nuclear import of MRTF hence the cytoplasmic accumulation of G-Actin leads to SRF transcription inhibition (Miralles *et al*, 2003). In that way, the Actin dynamic was described as a potent intracellular fulcrum for cell adaptation to its environment, connecting upstream RhoGTPases to downstream MRTF-SRF pathway (Hill *et al*, 1995; Sotiropoulos *et al*, 1999; Maekawa, 1999).

As mentioned above, studies on podocytes reported that CsA had anti-proteinuric effects via the stabilization of the Actin cytoskeleton structuring the foot processes. This stabilization relied on the overexpression and dephosphorylation of Cofilin. Even though the study did not focus on CFL oligomerization, the CFL: Actin ratio was in favour of CFL and the CFL concentration was higher, therefore the activity of CFL was regulated to promote Actin polymerization and stabilize microfilaments. What we observed in PTC was a destabilization of the Actin cytoskeleton structuring the intercellular convolutions. This indicated a different regulation of CFL activity where CFL would have to i) exert low saturation of F-Actin microfilaments (lower CFL: Actin ratio; lower levels of CFL oligomers), ii) be active for Actin-related functions i.e. dephosphorylated.

CsA had already been reported to elicit the remodelling of the Actin cytoskeleton of PTC (Martin-Martin *et al*, 2012). In this study, CsA (4.2 μM, 24 h) activated RhoA and the phosphorylation cascade, which led to CFL phosphorylation by LIMK. CFL inhibition results in the formation of stress fibres and the tightening of tight junctions. Our findings are just the opposite of these previous results and are incompatible with the previously described mechanism. Indeed, pharmacological inhibition of the RhoGTPases pathway elicited CsA like effects i.e. the loss of proximal tubular morphology and the inhibition of MRTF-SRF activity. These discrepancies might result from differences in cell culture conditions as, in the study by Martin-Martin et al, control LLC PK-1 seemed non-confluent, less differentiated, with non-convoluted membranes and a significant amount of stress fibres. Although the cell culture protocol appeared to be similar, the serum deprivation prior to drug exposure differed as it is not supplemented with hormones allowing advanced epithelial differentiation of LLC PK-1 cells at the morphological and cytoskeletal levels. Our present in vitro model seemed closer to the in vivo aspect of PTC than the in vitro one Martin-Martin et al developed.

Outside Actin-related functions, CFL is known to interact with Na^+^/K^+^-ATPase to provide energetic fuel to the pump motion. CFL binds triose phosphate isomerase and interacts with the alpha subunit of Na^+^/K^+^-ATPase to locally provide ATP. When Na^+^/K^+^-ATPase is inhibited (by potassium depletion or the action of a pharmacological inhibitor), a feedback mechanism disrupts energetic supply by CFL dephosphorylation (Jung *et al*, 2002, 2006a, 2006b).

CsA effects on TCP were reminiscent of OUA effects on HeLa cells (Jung *et al*, 2006a). OUA exerted a similar reorganization of the Actin cytoskeleton by dephosphorylating CFL via a mechanism where the inhibition of Na^+^/K^+^-ATPase activates the Ras/Raf/MEK cascade, in a Src-EGFR-dependent way, leading to the inhibition of LIMK downstream the small RhoGTPases pathway. OUA effects were reproduced by Y27632 just like CsA effects were mimicked by the RhoGTPases inhibitors.

Earlier findings about the mechanism of Na^+^/K^+^-ATPase energetic supply by pCFL-TPI complexes (Jung *et al*, 2002) reported that the overexpression of constitutively active CFL (Ser3 was replaced by Ala) was sufficient to cut endogenous CFL out of phosphorylation/dephosphorylation-based regulation which made TPI localization insensitive to the activation/inhibition of the RhoGTPases pathway. Likewise, in our study, the addition of S3-R was enough to partially exclude endogenous CFL from CsA-induced regulation.

Dephosphorylation of CFL by Slingshot phosphatases seemed unlikely since Slingshot phosphatases are activated by Calcineurin. CsA effects on CFL were unlikely to be related to Calcineurin since Tac and VIVIT did not elicit CsA-like effects on the Actin cytoskeleton or the MRTF-SRF transcription activity.

Numerous studies have already reported the inhibition of Na^+^/K^+^-ATPase by CsA, in vitro and in vivo. However, the observations were limited to the consequences in ion transport (Ihara *et al*, 1990; Tumlin & Sands, 1993; Lea *et al*, 1994; Ferrer-Martínez *et al*, 1996; Deppe *et al*, 1997; Younes-Ibrahim *et al*, 2003; Marakhova *et al*, 1998, 1999). Although this work is not the first one to study the relationship between the Na^+^/K^+^-ATPase pump and the Actin cytoskeleton, the partial inhibition (about 25%) of proximal tubular Na^+^/K^+^-ATPase by CsA has been the first observation of a direct correlation between variations in the pump activity and cytoskeletal remodelling so far. Besides, the potentiation of the CsA-induced depolymerization of F-Actin by OUA is in favour of a Na^+^/K^+^-ATPase -related mechanism.

How CsA inhibits Na^+^/K^+^-ATPase has yet to be elucidated. CsA is known to induce overexpression of endothelin, which is a well-known inhibitor of Na^+^/K^+^-ATPase activity (Zeidel *et al*, 1989; Nakahama, 1990). Furthermore, CsA can downregulate Na^+^/K^+^-ATPase activity through the inhibition of Cyclophilin B. The PPIase was described as a crucial partner for the structure and activity of Na^+^/K^+^-ATPase catalytic subunit (Suñé *et al*, 2010). Besides, the proteome monitoring of HEK cells by SILAC-LC-MS/MS reported the decrease in CypB protein abundance upon CsA exposure (Lamoureux *et al*, 2011). These findings, together with the observations of CsA-like effects upon CAI exposure, support the implication of Cyps in the inhibition of Na^+^/K^+^-ATPase by CsA. To what extent the Cyps are implicated has yet to be elucidated. However, our work suggests another mechanism leading to Na^+^/K^+^-ATPase inhibition.

Indeed, our molecular modelling of CsA docking into Na^+^/K^+^-ATPase was performed to visualise the potential interactions. CsA is mostly made of hydrophobic amino acids favouring its membrane partitioning (Haynes *et al*, 1985). Therefore, the preferential binding site to Na^+^/K^+^-ATPase is likely to be in the transmembrane site C, known to bind hydrophobic compounds such as lipids and cholesterol. Such binding is mostly expected to hinder proper motions of the transmembrane helices by impairing specific protein-lipid interactions. This may explain the experimentally observed inhibition of Na^+^/K^+^-ATPase by CsA given that the Na^+^/K^+^-ATPase (i) undergoes large-scale conformational changes during its reaction cycle and (ii) its activity is strongly dependent of the membrane environment (Bhatia *et al*, 2016). Interestingly, in the open structure, there are no binding sites in the transmembrane region, but there are two at the membrane interface that could (i) impair domain motions and (ii) prevent the proper closing of N– and A– domains during the reaction cycle. It is noteworthy that there is also a binding site under the A-domain, similar to the one in the closed conformation. Such preliminary results are in favour of direct interactions between CsA and Na^+^/K^+^-ATPase leading to impairment of large-scale conformational motions. It is important to note that the confirmation of the aforementioned molecular mechanism would require an important computational effort by biased molecular dynamics simulations, which if off-topic of the present study.

Altogether, these results let us propose a model (Figure 7) where: i) the inhibition of Na^+^/K^+^-ATPase by CsA leads to the dephosphorylation of CFL, independently from the classical regulation pathways; ii) CsA decreases the CFL: Actin ratio and the protein abundance of F-Actin-stabilizing and polymerization-promoting CFL oligomers; iii) active, minority, monomeric CFL depolymerizes low-saturated F-Actin microfilaments and replenish G-Actin pools; iv) the accumulation of G-Actin leads to the cytoplasmic trapping of the SRF co-factor MRTF; v) the low nuclear abundance of MRTF results in the low abundance of transcription-ready MRTF-SRF complexes and the decrease in MRTF-SRF transcription activity; overall, vi) CsA inhibits gene transcription by the MRTF-SRF complex via drug-specific, G-Actin-prone Actin-dynamics.

**Figure 7.**
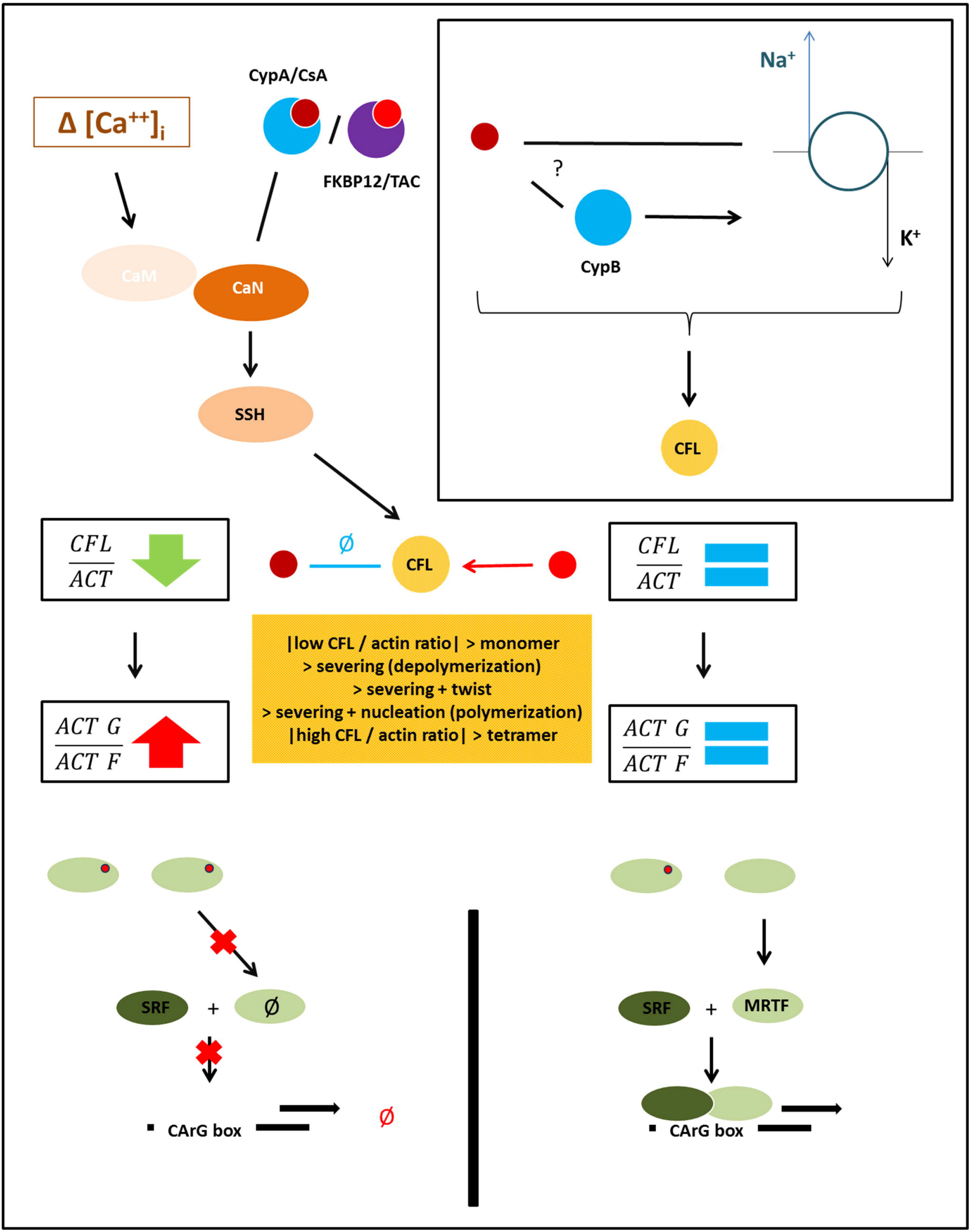
Proposed model of CsA-induced MRTF-SRF inhibition through Na^+^/K^+^-ATPase inhibition and remodelling of the Actin cytoskeleton.

The CsA-induced inhibition of MRTF-SRF gene transcription might have feedback consequences on Actin dynamics since ACTB, Cfl1 and other ABP-coding genes are identified genes of the CArGome (the ensemble of MRTF-SRF genes whose promoters contain CArG boxes) and the activity of the MRTF-SRF complex was shown to be required for the homeostasis of Actin levels and the organization of the Actin cytoskeleton (Sun, 2005).

As a matter of facts, cellular consequences of MRTF-SRF inhibition might be large-scale perturbations since MRTF-SRF regulates the transcription of genes coding for transport and adhesion proteins, enzymes of the lipid and glucose metabolisms, transcription factors, growth factors, growth factor receptors, etc.

For instance, past studies clearly elucidated the role of MRTF-SRF inhibition in multifactorial biological processes like proapoptotic mechanisms (Cao *et al*, 2011; Sisson *et al*, 2015) and epithelial-mesenchymal transition (Korol *et al*, 2016; Gasparics & Sebe, 2018), which have been widely described as CsA-related pathophysiological mechanisms.

Hence why the proposed model of an inhibition of MRTF-SRF transcription activity downstream an original regulation of Actin dynamics upon CsA exposure might be of utmost interest to explain known in vitro effects of CsA and provide a unified mechanism of CsA toxicity in PTC.

## Materials and methods

### Chemicals

Dulbecco’s Modified Eagle’s Medium-Ham’s F12 (DMEM-F12 1:1, #31331, Gibco), Fetal Bovine Serum (FBS, #10500), 1 M HEPES (#15630), 7.5 % Sodium bicarbonate (#25080), 10,0 UI.mL-1 Penicillin / Streptomycin (#15140), 0.05% Trypsin-EDTA (#25300-054) Dulbecco’s Phosphate Buffer Saline (PBS, #14190) were purchased from Gibco. Sodium selenite (#S5261), CsA (#30024), Tac (#F4679), Jasplakinolide (#J4580), Latrunculin A (#L5163), ROCK inhibitor Y27632 (#Y0503), Rac1 inhibitor EHT 1864 (#E1657), Alamethicin (#A4665), Digitonin (#D141), ATP (#A1852) Na+/K+-ATPase inhibitor Ouabain (#O3125), insulin (#I4011), triiodothyronine (#T6397), dexamethasone (#D4902), Protease Inhibitor Cocktail (#P8340) and Phosphatase Inhibitor Cocktail 2 (#P5726) were purchased from Sigma-Aldrich. Cyclophilin A inhibitor (#239836, Calbiochem) was purchased from Millipore. LIMK1/2 inhibitor LIMKi3 (#4745) was provided by Tocris.

S3-R peptide (MASGVMVSDDVVKVFNRRRRRRRR), an analogue of the Cofilin phosphorylation site, was synthetized by ProteoGenix (Schiltigheim, France).

### Cell culture conditions

LLC PK-1 (Lilly Laboratories Porcine Kidney-1) porcine proximal tubule cells (ATCC-CL-101, ATCC, Manassas, VA) were expanded in 75 cm^2^ flasks at 37 °C with 5 % CO2 and passed once confluence was reached. Culture medium consisted in a 1:1 DMEM-F12 mix supplemented with 5 % FBS, 15 mM HEPES, 0.1 % Sodium bicarbonate, 100 UI.mL-1 Penicillin / Streptomycin and 50 nM Sodium selenite. Passages were performed with 0.05% Trypsin-EDTA. LLC PK-1 cells were cultured between passage 7 and passage 30.

Seeded LLC PK-1 sustained serum starvation and were fed with hormonally-defined (25 μg.mL-1 insulin, 11 μg.mL-1 transferrin, 50 nM triiodothyronine, 0.1 μM dexamethasone, 0.1 μg.mL-1 desmopressin) fresh medium to engage epithelial differentiation, for 24 hours.

Hormonally-defined LLC PK-1 cells sustained cell transfection or were exposed for 24 hours to: vehicle mix (Ethanol, DMSO), 5 μM CsA, 0.05 μM Tac, 1 μM VIVIT (a specific NFAT inhibitor), 0.5 μM Cyclophilin A inhibitor, 200 nM Jasplakinolide, 100 nM Latrunculin A, 10 μM ROCK inhibitor Y27632, Rac1 inhibitor EHT 1864, and LIMK inhibitor LIMKi3, 100 nM Na+/K+-ATPase inhibitor Ouabain or 100 μg.L-1 S3-R.

### Actin cytoskeleton characterization

#### immunocytochemistry

Serum-starved, hormonally-defined, treated LLC PK-1 cells seeded on glass cover slips in 6-well plates were washed twice for 3 min in PBS. Then, cells were fixed in 4 % paraformaldehyde-PBS for 10 min, at room temperature. Cells were washed once again in PBS for 3 min then permeabilised with 0.1 % Triton/X-100-PBS for 5 min, at room temperature. Permeabilised cells were washed three times for 3 min in PBS before incubation in Phalloidin-TRITC-PBS for 30 min, at 37 °C, in the dark. Eventually, cover slips were washed three times for 3 min in PBS then mounted on glass microscope slides using ProLong^®^ Gold Antifade Reagent as a mounting medium and a DAPI nuclei staining. Florescence labelling was visualized using a Leica DM-RX microscope (16×40).

#### Image analysis

For each exposure condition, ten photographs (*.tiff) were randomly taken to report from an independent experiment. Image analysis was conducted using the ImageJ software (v.1.48). Early image processing consisted in a restriction to the green channel of the RGB picture. The implemented Auto Local Threshold tool further selected the fluorescence-positive pixels (Niblack thresholding method, radius = 15). Thus, for each photograph, a semi-quantitative ratio was calculated as a percentage of the total image area.

All above-mentioned steps were automatized thanks to the macro editing tool of the ImageJ software.

Statistical analysis of fluorescence area data was performed using the one-way analysis of variance (1-way ANOVA) test followed by the Bonferroni’s Multiple Comparison post-test with a significance threshold at p<0.05, as implemented in GraphPad Prism (v. 5.04).

### Luciferase Gene Reporter Assay for SRF/SRE Activity

#### Stable cell transfection & clone selection

Volumes equivalent to 0 μg, 2 μg or 4 μg of plasmid DNA (pGL4.34[luc2P/SRF-RE/Hygro] Vector, Promega) and transfection reagent lipofectamine (1:25, Lipofectamine™ 2000 Transfection Reagent, Invitrogen) were separately diluted in FBS-free, antibiotic-free routine medium. Then, DNA and lipofectamine preparations were blended and put at rest for 15 min, at room temperature. Serum-starved, hormonally-defined LLC PK-1 cells in 6-well plates were incubated with DNA-lipofectamine mixtures for 24 h, at 37 °C.

Transfected LLC PK-1 cells (LLC PK-1 SRE) were incubated in routine medium for 72 h before antibiotic selection to allow proper plasmid integration. Then, cells were re-suspended in 0.05 % Trypsin-EDTA, 10 min at 37 °C. Cell suspensions were diluted 10 times and reseeded in routine medium completed with 1 % Hygromycin B (50 mg.mL-1) refreshed every 48 h until colonies appear. Colonies were harvested thanks to trypsin-soaked paper disks (#Z374431, Sigma-Aldrich) and re-seeded to start routine culture in antibiotic-supplemented growth medium.

#### Luciferase gene reporter assay/Bioluminescence assay

Serum-starved, hormonally-defined, treated LLC PK-1 SRE were re-suspended mechanically in 1X lysis buffer (Cell Culture Lysis 5X Reagent, #E153A, Promega). Cell lysates were distributed in technical duplicates in a white 96-well plate (#3912, Costar). Bioluminescence signals were detected by the Enspire Multimode Reader (Perkin-Elmer).

### Oligomer cross-linking and Western blot

Serum-starved, hormonally-defined, treated LLC PK-1 cells in 60 mm Petri dishes were incubated in hormonally-defined medium – 1 % formaldehyde (cross-linking range 2.9 Å) for 15 min under agitation. Formaldehyde cross-linking was quenched by the addition of 1 M glycine (final concentration 125 mM). LLC PK-1 were washed twice with PBS and lysed by scrapping in a custom RIPA lysis buffer (150 mM NaCl, 50 mM TRIS-HCl, 0.1 % NP-40, 0.1 % SDS, 1 mM EDTA in ultrapure H_2_O, supplemented with a 1:100 anti-protease / antiphosphatase mix). Cell lysates were incubated on ice for 30 min and centrifuged for 15 min at 21000 g. Supernatants were stored until protein concentration was measured using the Bradford colorimetric method. Forty micrograms of proteins per exposure condition were separated by electrophoresis under reducing and denaturing conditions on a NuPAGE^®^ 12% Bis-Tris pre-cast gel (#NP0341, ThermoFisher) in 1X NuPAGE™ MOPS SDS running buffer (#NP0001, ThermoFisher) and transferred to a nitrocellulose (NC) membrane (NP23001, ThermoFisher) using the iBLOT 2 Dry Blotting system (#IB21001, ThermoFisher). Membranes were stained with 0.1 % Ponceau S, 5 % acetic acid solution for total protein visualisation. Membranes were blocked for 1 h at room temperature under agitation with TBS-Tween (TBS-T) buffer (10 mM Tris 7.6, 150 mM NaCl, 0.1 % Tween-20) complemented with 5 % (W/V) non-fat milk powder to obtain BLOTTO buffer. Primary antibody incubation (anti-CFL, 1:10,000, #PA1-24931, ThermoFisher) was done in BLOTTO for 1 h at room temperature. After three 5-min washes in TBS-T, secondary antibody incubation (F(ab’)2-Goat anti-Rabbit IgG (H+L) Secondary Antibody, HRP, 1:10,000, #A24531, ThermoFisher) was performed in BLOTTO for 1 h at room temperature then washed again. Membranes were incubated in a 1: 1 mix of SuperSignal™ West Pico PLUS Chemiluminescent Substrate kit (#34577, ThermoFisher) and analyzed by the ChemiDoc Imaging system (Bio-Rad) for chemiluminescent signal detection and acquisition. Quantitation was computed via the ImageLab software (Bio-Rad) after total protein normalisation.

### Na^+^/K^+^-ATPase activity/Phosphate release assay

Serum-starved, hormonally-defined, treated LLC PK-1 cells in 24-well plates were permeabilised by osmotic shock in ultrapure water (150 μL) and fast freeze in liquid nitrogen (10 s). Cells were thawed (200 μL) in a buffer with a composition of 200 mM sodium chloride, 80 mM histidine, 20 mM potassium chloride, 6 mM magnesium chloride, 2 mm EGTA, 2 μg.mL-1 Alamethicin, 30 μM Digitonin. For each drug exposure condition, a volume of 30 mM Ouabain (final concentration 1 mM) or ultrapure water was added to every other well. Cells were either incubated 30 min at 37 °C. A volume of 30 mM ATP (final concentration 1 mM) was added to each well. Cells were incubated 30 min more at 37 °C. On ice, trichloroacetic acid: water (1:1) terminated ATP hydrolysis reactions. Plates were centrifuged at 3000 rpm for 10 min at room temperature. Supernatants were diluted 50 times and distributed (50 μL) in technical triplicates to a 96-well plate.

Concentration of free phosphate released from ATP hydrolysis was measured using the BIOMOL^®^ Green kit (BML-AK-111-0250, Enzo). Na^+^/K^+^-ATPase activity (expressed in nmol of released phosphate/30min/105 cells) was calculated as the difference in free phosphate concentration in the presence and absence of Ouabain.

### iTRAQ shotgun proteomics

#### Protein extraction, sample preparation, iTRAQ labelling and isoelectric focusing

After 24 h drug exposure, LLC PK-1 cells were washed twice with PBS and lysed by scrapping in a custom RIPA lysis buffer (150 mM NaCl, 50 mM TRIS-HCl, 0.1 % NP-40, 0.1 % SDS, 1 mM EDTA in ultrapure H_2_O, supplemented with an anti-protease / antiphosphatase mix). Cell lysates were incubated on ice for 30 min and centrifuged for 15 min at 21000 g. Supernatants were stored until protein concentration was measured using the Bradford colorimetric method and iTRAQ labeling. Twenty-five micrograms of proteins were precipitated by – 20 °C cold acetone. After acetone evaporation, the precipitates were solubilized in 25 mM ammonium bicarbonate then were incubated with 50 mM dithiothreitol for 40 min at 60 °C, to reduce disulfide bonds, 100 mM iodoacetamide in the dark for 40 min at room temperature, to alkylate/block cysteine residues and eventually were digested for 24 h at 37 °C with mass-spectrometry grade trypsin (#V5280, Promega) at a 1: 50 enzyme: substrate ratio.

After digestion, samples were incubated with iTRAQ tags (iTRAQ Reagents Multi-plex kits, 4-plex, #4352135, Sigma-Aldrich) – one tag per drug exposure condition and five different tag/condition associations over five experiments – for 1 h at room temperature. Labeled samples were mixed and separated into 12 fractions by isoelectric focusing (OFFGEL 3100 Fractionator, Agilent Technologies, Santa Clara, CA) for 24 h at increasing voltage and steady intensity of 50 μA in a 3-10 pH IPG strip. Fractions were retrieved for further MS analysis after the IPG strip was incubated in a 1:1 acetonitrile (ACN): water, 0.1 % formic acid (FA) wash solution for 15 min at room temperature.

#### nano-LC peptide separation and Q-Q-TOF mass spectrometry

IEF fractions were analyzed by nano-LC MS/MS using a nano-chromatography liquid Ultimate 3000 system (LC Packings DIONEX, Sunnyvale, CA) coupled to a Triple TOF 5600+ mass spectrometer (ABSciex, Toronto, Canada). For each sample, 5 μL were injected into a pre-column (C18 Pepmap™ 300 μm ID × 5 mm, LC Packings DIONEX) using the loading unit. After desalting for 3 min with loading solvent (2 % ACN, 0.05 % trifluoroacetic acid (TFA)), the pre-column was switched online with the analytical column (C18 Pepmap™ 75 μm ID × 150 mm, LC Packings DIONEX) pre-equilibrated with 95 % solvent A (ACN 5 % – FA 0.1 %). Peptides were eluted from the pre-column into the analytical column and then into the mass spectrometer by a linear gradient from 5 % to 25 % in 70 min, then to 95 % of solvent B (98 % ACN, 0.1 % FA) over 120 min at a flow rate of 200 nL/min.

Data acquisition was carried out by IDA (Information-Dependent Acquisition) mode of Analyst 1.7 TF software (ABSciex). The data from MS and MS/MS were continuously recorded with a cyclic duration of 3 s. For each MS scan, up to 50 precursors were selected for fragmentation based on their intensity (greater than 20,000 cps), their charge state (2+, 3+) and if the precursor had already been selected for fragmentation (dynamic exclusion). The collision energies were automatically adjusted according to charge state, ionic mass of selected precursors and iTRAQ labelling.

#### Mass spectrometry data processing and relative protein identification / quantification

MS and MS/MS data for five independent experiments (biological replicates) (*.wiff, 1 per fraction, 12 files per experiment) were submitted to Mascot Server 2.2.03 via ProteinPilot (version 5.0, ABSciex) for protein identification, and searched against two complementary *Sus scrofa* databases: a Swiss-Prot database (2015_10 release) and a TrEMBL database (2015_10 release). Carbamidomethyl (C) was defined as a fixed modification. Oxidation (O), iTRAQ4plex (K), iTRAQ4plex (Y), iTRAQ4plex (N-term) were defined as variable modifications. MS/MS fragment mass tolerance was set at 0.3 Da. Precursor mass tolerance was set at 0.2 Da.

Mascot raw data files (*.dat, 1 per experiment) were saved for further isobaric tags-based peptide and protein quantitation with the Java implementation of the Quant algorithm, jTRAQx (version 1.13, (Muth *et al*, 2010)). Reporter mass tolerance was set at 0.05 Da while iTRAQ correction factors were implemented as provided by ABSciex. This tool generated one *.jpf file (tab-delimited text file) for each series.

*.jpf were submitted to the CiR-C (Customizable iTRAQ Ratio Calculator) algorithm which excluded irrelevant data according to: i) identification confidence: peptides are retained if the probability that the observed positive match is a random match is below 5 % (p < 0.05, Mascot score > 30); ii) quantification confidence: peptides are retained if all iTRAQ ratios have been successfully calculated i.e., peptides with 0.0 ratios or uncalculated ratios (null ratios) are discarded; iii) peptides related to ‘Fragment’– and ‘REVERSED’-annotated proteins are discarded. After irrelevant data removal, CiR-C drew up an exhaustive catalogue of identified peptide sequences with their associated Swiss-Prot or TrEMBL accession IDs. Peptides were assigned to a frequency index of positive matches (identification in {1;2;3;4;5} out of 5 independent experiments) and CiR-C drew a second catalogue of peptides with the highest frequency index (n=5). Protein ratios were calculated as both overall and series-specific median values of peptide ratios associated with a given accession ID and frequency index.

#### PANTHER Overrepresentation test

The protein list analysis was performed by submitting Swiss-Prot accession IDs to the online tool powered by the PANTHER Classification system. The PANTHER Overrepresentation test (release date 20170413) parsed the PANTHER database (version 12.0, released on 2017-07-10) using the *Sus scrofa* reference list and the PANTHER Protein Class annotation data set. Only p<0.05 items were retained and considered significantly over-represented.

#### Visualisation of protein networks

The visualisation of protein networks was performed by submitting Swiss-Prot accession IDs to the online tool STRING.

#### Biological significance criteria

Frequency distribution of the iTRAQ peak intensities ratios could be approximated as a Gaussian distribution: iTRAQ peak intensities ratios frequency distribution significantly fitted implemented Gaussian non-linear regression (n = 370, R^2^ = 0.9946, mean = 1.02 ± 0.10). One of the Gaussian distribution properties is: I) 66% of the values lie in a 1 S.D. range around the mean, II) 95 % of the values lie in a 2 S.D. range around the mean, III) 99 % of the values lie in a 3 S.D. range around the mean. Applying the “mean ± n S.D.” property to the set of iTRAQ peak intensity ratios, the upper thresholds for significant upregulations upon drug exposure would be “1.02 – 0.10 = 0.92”, “1.02 – 2 × 0.10 = 0.82” and “1.02 – 3 × 0.10 = 0.72” while the lower thresholds for significant downregulations upon drug exposure would be “1.02 + 0.10 = 1.12”, “1.02 + 2 × 0.10 = 1.22” and “1.02 + 3 × 0.10 = 1.32”. Thus, proteins were classified thanks to their distance from the unity: proteins with median ratios between 0.92 and 1.12 were annotated as non-impacted proteins; up-regulated proteins with quantitative ratios beyond 1.12 were ranked according to three tiers of significant fold increase: +σ (1.12 ≤ r < 1.22), +2σ (1.22 ≤ r < 1.32) or +3σ (1.32 ≤ r). The same way down-regulated proteins with quantitative ratios below 0.82 were ranked according to three tiers of significant fold decrease: −σ (0.92 ≥ r > 0.82), −2σ (0.82 ≥ r > 0.72) or −3σ (0.72 ≥ r).

#### Hierarchical clustering & heat-map representation

Heat map representation and hierarchical clustering were generated using the Euclidian distance calculation as provided by the GENE-E software (version 3.0.204).

#### Data availability

The MS proteomics data have been deposited to the ProteomeXchange Consortium (http://proteomecentral.proteomexchange.org) (Deutsch *et al*, 2017) via the PRIDE partner repository (Vizcaíno *et al*, 2016) with the data set identifier PXD007891 (username, password: TPSGICw9).

### Setup of cyclosporine docking into Na^+^/K^+^-ATPase

A cyclosporine structure was docked into a human-sequence homology model of Na^+^/K^+^-ATPase based on crystal structures (PDB IDs: 2ZXE and 4HQJ respectively for the open conformation E2 state (Shinoda *et al*, 2009), and the closed conformation E1 state (Nyblom *et al*, 2013)).

The molecules were prepared for docking in Autodock Tools (Morris *et al*, 2009) and the docking was performed in Autodock Vina (Trott & Olson, 2009), with grid covering the whole protein and exhaustiveness set to 400. A second docking was performed into the lipid binding site C to better evaluate the cyclosporine binding in this position.

The docking was performed in three variations, with 20 binding poses generated for each case: i) the closed structure E1 with a bound ATP molecule, ii) the closed structure E1 without ATP, iii) the open structure E2 without phosphate.

Given the similarity of the docking results between the closed conformation with and without ATP, and the experimental conditions, only the results of the docking with ATP are presented. The residues taking part in ligand binding were evaluated using PLIP (Salentin *et al*, 2015). Figures were made using PyMOL (The PyMOL Molecular Graphics System, Version 1.6, Schrödinger, LLC).

## Acknowledgements

This research was partially supported by the grant No. LO1204 (Sustainable development of research in the Centre of the Region Haná) from the National Program of Sustainability I, MEYS.

## Author contributions

BB and ME planned and designed the study. BB designed, performed and analysed cell biology and molecular biology experiments with HA, QF (cofilin oligomer cross-linking and Western blotting / Na^+^/K^+^-ATPase activity measurement) and FLS (iTRAQ shotgun proteomics). PC and FDM designed, performed and analysed molecular modelling experiments. BB wrote the manuscript. All authors reviewed and edited the manuscript.

## Expanded View figure legends

Figure EV1 – Dynamic mapping of LLC PK-1 proteome highlights CNI-specific expression profiles of Actin family cytoskeletal proteins.

A. STRING visualization of iTRAQ-monitored Actin family cytoskeletal protein network (PANTHER Protein Class PC00041)
B. Heat-map representation of the identified Actin family cytoskeletal proteins. Cut-offs for biological significance were calculated as Mean ± 1 SD (1.02 ± 0.10) based on the approximation of the iTRAQ ratio frequency distribution. Lower cut-off: 0.92 (green); upper cut-off: 1.12 (red).

Figure EV2 – Inhibitors of the RhoGTPases pathway elicited CsA-like features and potentiated CsA effects

A. Quantification of red fluorescence-positive area. Mean ± SEM, One-way ANOVA plus Tukey’s post-test (p<0.01**,p<0.001***) (n=3).
B. SRF transcription activity was measured by luciferase gene reporter assay in LLC PK-1 SRE. Mean ± SEM. One-sample t-test for versus control comparison, One-way ANOVA plus Tukey’s post-test for multiple condition comparison (p<0.01**,p<0.001***) (n=3)

Drug condition: Vehicle (0.5% Ethanol-0,2% DMSO), 5 μM CsA, 10 μM Y27632, Y27632 + CsA, 10 μM EHT1864, EHT1864 + CsA. Exposure time: 24 h.

Figure EV3 – Molecular modelling of CsA docking into Na^+^/K^+^-ATPase

A. Binding poses of CsA docking into Na^+^/K^+-^ATPase open conformation (E2 state)
B. Binding poses of CsA docking into Na^+^/K^+^-ATPase closed conformation (E1 state)
C. Close-up of site C-located binding poses of CsA docking into Na^+^/K^+^-ATPase closed conformation (E1 state)

