## Supplementary figures and images for "Cyclosporine A inhibits MRTF-SRF signalling through Na^+^/K^+^ ATPase inhibition & Actin remodelling"

### Figure EV1

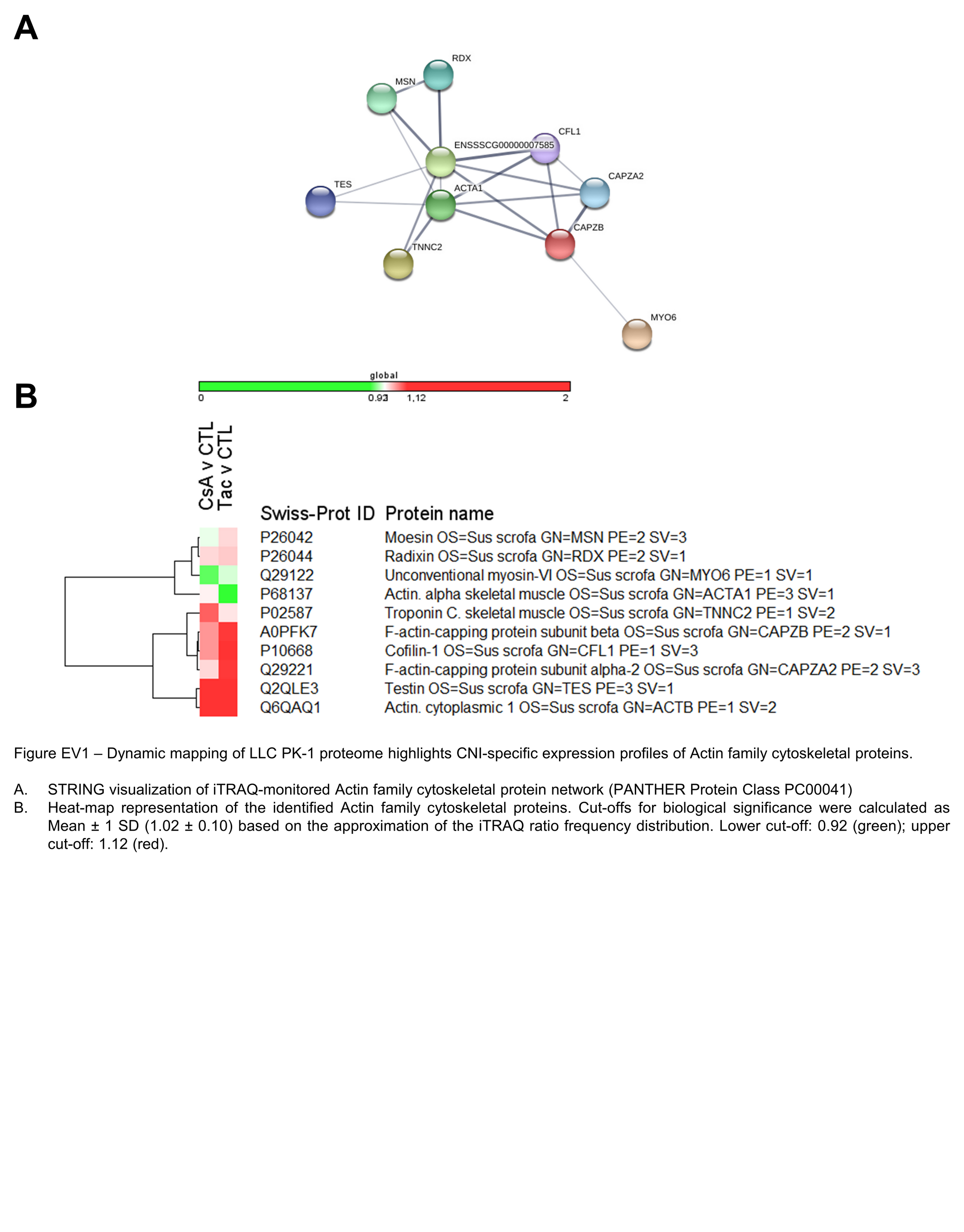

### Figure EV2

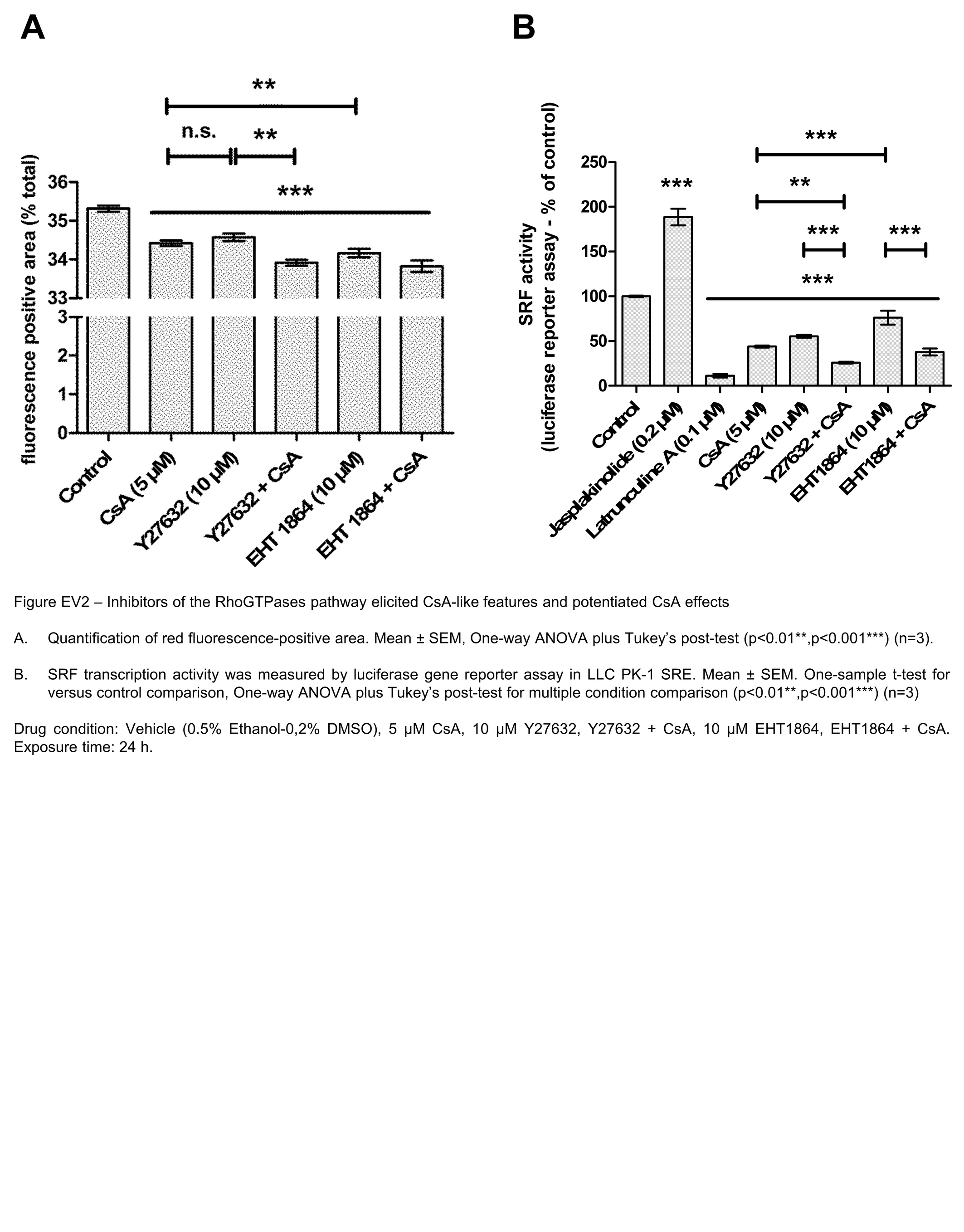

### Figure EV3

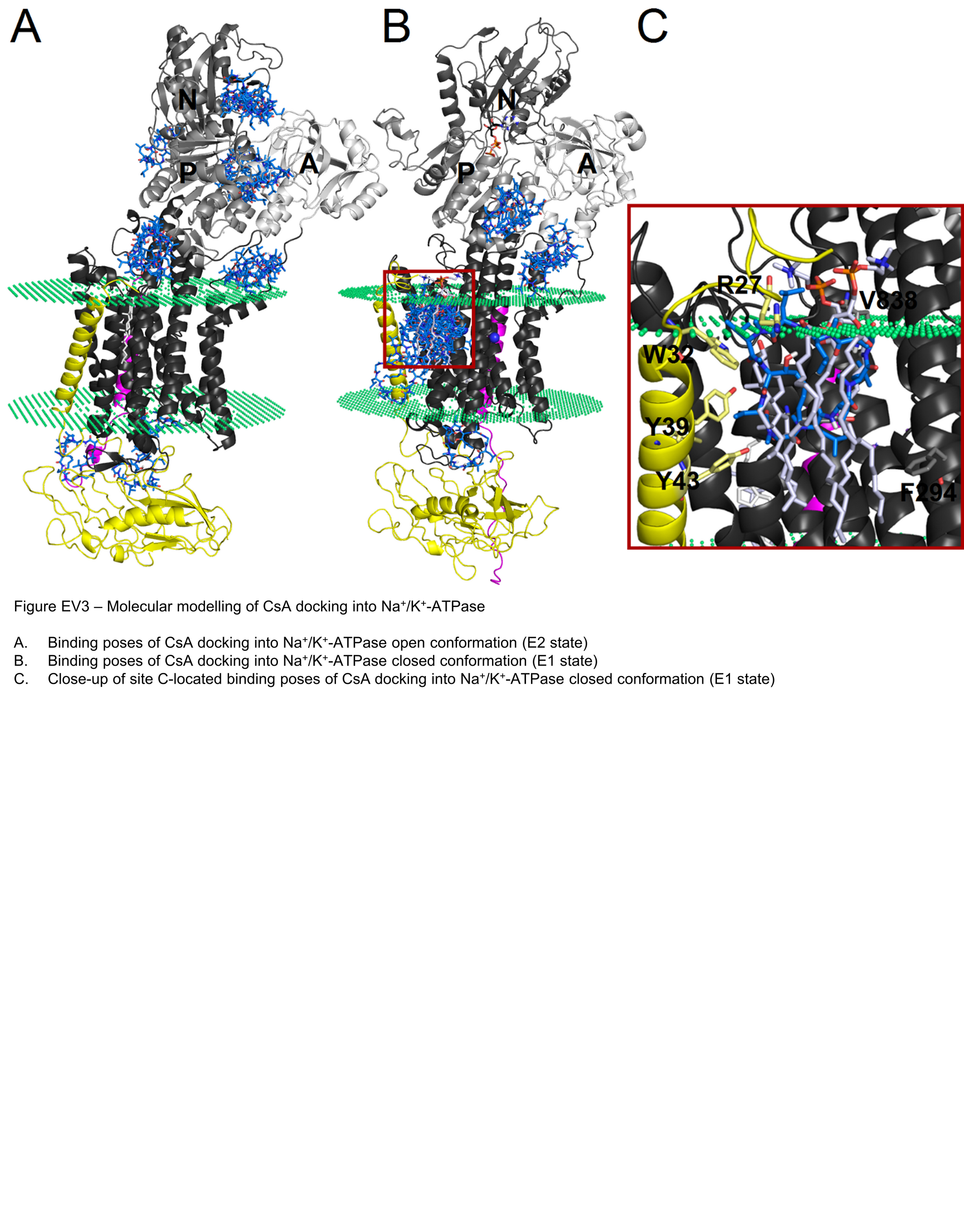
